# High-throughput proteomic and phosphoproteomic analysis of formalin-fixed paraffin-embedded tissue

**DOI:** 10.1101/2024.11.17.624038

**Authors:** Moe Haines, John R. Thorup, Simone Gohsman, Claudia Ctortecka, Chelsea Newton, Dan C. Rohrer, Galen Hostetter, D. R. Mani, Michael A. Gillette, Shankha Satpathy, Steven A. Carr

## Abstract

Formalin-fixed, paraffin-embedded (FFPE) patient tissues are a valuable resource for proteomic studies with the potential to associate the derived molecular insights with clinical annotations and outcomes. Here we present an optimized, partially automated workflow for FFPE proteomics combining pathology-guided macro-dissection, Adaptive Focused Acoustics (AFA) sonication for lysis and decrosslinking, S-Trap digestion to peptides, and liquid chromatography-tandem mass spectrometry (LC-MS/MS) analysis using Orbitrap, Astral or timsTOF HT instrumentation. The workflow enables analysis of up to 96 dissected FFPE tissue samples or 10 µm scrolls, identifying 8,000-10,000 unique proteins per sample with median CVs <20%. Key optimizations include improved tissue lysis strategies, protein quantification for normalization, and peptide cleanup prior to LC-MS/MS analysis. Application to lung adenocarcinoma (LUAD) FFPE blocks confirms the platform’s effectiveness in processing complex, clinically relevant samples, achieving deep proteome coverage and quantitative robustness comparable to TMT-based methods. Using the newly released Orbitrap Astral with short, 24-minute gradients, the workflow identifies up to ∼10,000 unique proteins and ∼11,000 localized phosphosites in LUAD FFPE tissue. This high-throughput, scalable workflow advances biomarker discovery and proteomic research in archival tissue samples.

## Introduction

Formalin-fixed, paraffin-embedded (FFPE) tissue samples with clinical annotation represent an invaluable resource. Patient tissue profiling by global proteomics using liquid chromatography-tandem mass spectrometry (LC-MS/MS) provides a molecular foundation for exploration of disease mechanisms and prediction of clinical outcomes that goes beyond what can be achieved by traditional pathology or immunohistochemistry (1–5). The extensive collections of FFPE samples in biobanks and pathology archives facilitate large-scale studies, aiding in biological discovery and the identification of candidate biomarkers and molecular alterations associated with diverse pathologies (6).

Mass spectrometry-based proteomic analyses of FFPE tissues present a number of challenges. Many FFPE proteomics workflows have relied on laborious xylene and ethanol washes for deparaffinization (7, 8), although alternative approaches have started to emerge such as the use of the Beatbox (Preomics), which leverages bead-based tissue lysis (9); substitution of SafeClear (Fisher Scientific) for xylene (10); and lysis using the Covaris Adaptive Focused Acoustics (AFA) technology (3, 11–13). Despite these innovations, extraction of proteins and peptides from FFPE tissues has proven difficult, constraining depth of coverage and quantitative characterization of post-translational modifications (PTMs) (4, 7, 14). Tissue preservation techniques can also impact quantitative reproducibility relative to fresh frozen tissue (15) and contaminants such as wax in FFPE can be problematic for downstream nanoflow LC-MS/MS applications. Process improvements are needed to achieve deep-scale analysis of FFPE tissues and take full advantage of the invaluable resource afforded by repositories of well-annotated FFPE specimens.

In this study, we describe a highly optimized and partially automated workflow that enables reproducible deepscale proteomic analysis of FFPE tissue yielding ∼8,000-10,000 unique proteins and up to ∼14,000 localized phosphosites per sample. The plate-based workflow employs the Covaris AFA sonicator for FFPE sample processing and decrosslinking, an Opentrons OT-2 robot for complex liquid handling steps, positive pressure manifold-assisted S-Trap-based digestion and purification, and Data Independent Acquisition (DIA) analysis on Orbitrap or timsTOF instrumentation. Additionally, DIA data was uploaded to Terra (https://app.terra.bio/), a cloud-based environment running on Linux that is designed to support Spectronaut (16), FragPipe (17), and DIA-NN (18), enabling fast and scalable data processing to meet the intensive computational demands of DIA pipelines in large-scale experiments. The set-up allows us to process and analyze up to 96 samples in less than 4.5 days.

## Experimental procedure

### Deparaffinization of FFPE Tissue using Covaris Ultrasonicator

Breast (BRC) and colorectal cancer (CRC) FFPE blocks (∼2 x 3 cm tissue area), provided by the Van Andel Research Institute (VARI), were sectioned into serial 10 µm scrolls using the HistoCore MULTICUT (Leica). Scrolls were transferred immediately into a Covaris 96 AFA-Tube TPX Plate (Covaris, Part No. 520291) and fully submerged in wells prefilled with Covaris tissue lysis buffer (TLB) (Covaris, Part No. 52-284). FFPE tissue was deparaffinized by ultra-sonication in the Covaris LE220Rsc Focused-ultrasonicator at 3 mm Y-dither at 10 mm/s, followed by heating to 90°C in a thermocycler (Eppendorf) with a heated lid at 105°C for 90 minutes to de-crosslink proteins. The tissue was then sonicated again for homogenization at 1 mm Z-dither at 20 mm/s.

For deparaffinization and homogenization, the ultrasonicator was maintained at 20°C, operated with a peak power of 350 W, and set to 200 cycles per burst; 5 minutes of sonication was applied to each lane of the plate. The parameter adjustments evaluated included increasing the Covaris default sonication duty factor (DF) from 25% to 50% and using either 100 µL or 150 µL of TLB. Optimal sonication conditions of 50% in 150 µL of TLB were used for subsequent experiments. Total protein amount following sonication was quantified using a BCA protein assay kit (Thermo Fisher, Cat No. 23225), with standards diluted 5X in TLB. The Covaris plate was spun down briefly at 1500 *g* in a centrifuge (Beckman Coulter) and 2 µL of the semi-clarified lysate was diluted 5X in water for quantification.

### Sample cleanup and digestion optimization

For protein digestion and cleanup, suspension trapping (S-Trap) and solvent precipitation bead (SP3) workflows were evaluated. All lysates were initially treated with 2 mM magnesium chloride and 0.8 U/µL of benzonuclease (Sigma Aldrich, Cat No. E8263-5KU). BRC and CRC samples were normalized to 50 µg of protein lysate, followed by reduction and alkylation in 5 mM tris(2 - carboxyethyl)phosphine (TCEP) (Thermo Fisher, Cat No. 77720) and 50 mM 2-Chloroacetamide (CAA) (VWR, Cat No. TCC2536-25G) for one hour while mixing in darkness at room temperature (RT).

For the SP3 workflow, the beads (Cytiva Sera-Mag Hydrophilic and Sera-Mag Hydrophobic Magnetic Particles: Cat No. 44152105050250 and 24152105050250) were equilibrated at RT for 30 minutes, mixed at a 1:1 ratio, washed three times with 80% acetonitrile (MeCN), and resuspended at 50 mg/mL. The bead mix was spiked into each sample at a bead to protein ratio of 10:1. Protein binding to the beads was initiated by adding MeCN to a final concentration of 80%. The beads were mixed during binding for 5 minutes and pelleted using a magnetic rack, prior to removal of the supernatant by manual aspiration. The beads were then washed three times with 500 µL of 80% MeCN. On-bead protein digestion was performed overnight in 50 mM triethylammonium bicarbonate (TEAB) pH 9 with mass spectrometry-grade trypsin (Promega) and LysC (FUJIFILM) at a ratio of 1:50 (weight/weight). Following overnight digestion, the beads were centrifuged at 3000 *g* and the supernatant was collected. The beads were then washed with 50 µL of water followed by 1% FA; in each wash step, the samples were centrifuged and transferred to a magnetic rack, where the supernatants were collected and combined.

For suspension trapping, 1.2% phosphoric acid was added to each reduced and alkylated sample, followed by six times the volume of 100 mM TEAB in 90% methanol (MeOH) (pH 7.1). Samples were mixed thoroughly via repeated pipette aspirations and loaded on the S-trap^TM^ 96-well Mini Plate (Protifi) using a vacuum manifold. After loading, the wells were washed three times with 300 µL of a 50/50 (volume/volume) chloroform/methanol solution and three times with 300 µL of 100 mM TEAB in 90% MeOH pH 7.1. Samples were digested overnight in 125 µL of 50 mM TEAB pH 9, using trypsin and LysC at a ratio of 1:50 (weight/weight). Peptides were eluted into a deep-well plate (Cytiva, Part No. 7701-5200) with sequential 80 µL additions of 50 mM TEAB pH 9, 0.2% formic acid (FA), and 0.2% FA in 50% MeCN.

### StageTip desalting optimization

Peptide digests were quantified using a quantitative fluorometric peptide assay (Thermo Fisher, Cat No. 23290), acidified with 1% FA, and a specified amount StageTip-desalted with Empore C18 disks (CDS Analytical, Part No. 98-0604-0217-3). The StageTips were conditioned with 100% MeOH and 0.1% FA in 50% MeCN, then equilibrated with 1% FA. Peptides were loaded, washed three times with 0.1% FA, and eluted with 0.1% FA in 50% MeCN. Samples were dried down in a speedvac vacuum concentrator and stored at −80°C until LC-MS/MS analysis.

SDB-RPS StageTips (Empore, Cat No. ST7600196) were implemented to reduce build-up on the column emitter. Peptides were acidified with 1% FA and StageTips conditioned with 100% MeCN and then equilibrated with 1% FA before loading. The peptides were washed twice with 1% FA in isopropyl alcohol (IPA), followed by 1% FA. Peptides were eluted with 5% ammonium hydroxide (NH_3_OH) in 80% MeCN. The samples were dried down and stored at −80°C until LC-MS analysis.

SDB-XC StageTips were prepared by packing SDB-XC (Empore, Cat. No 98060402231EA) punches at a ratio of 1 punch for each 1 ug of peptides. The SDB-XC StageTip was conditioned with 100% of MeOH and 0.1% FA in 50% MeCN, then equilibrated with 1% FA. Peptides were acidified to 1% FA and loaded on the conditioned SDB-XC StageTip for fractionation. The StageTip was washed twice with 1% FA in water. Peptides were eluted from the StageTip in five steps with 0.1% NH3OH in 5%, 10%, 15%, 20%, and 80% MeCN. Samples were dried and stored at −80°C until LC-MS analysis.

### Semi-automated 96-well plate workflow

Automation was implemented to enable efficient 96-well plate processing of FFPE tissue. Following plate-based Covaris sonication and S-trap loading of full pellets, the requisite trypsin and Lys-C quantities were determined based on per-sample protein yield. An Opentrons OT-2 Robot was used to prepare the digestion enzymes using a custom Python script to read a CSV file specifying the volume of protease for each sample in a 96-well plate before adjusting each solution to a final volume of 100 µL with 50 mM TEAB. The script and CSV file were uploaded into the Opentrons application programming interface (API) for variable input steps. A multi-channel pipette was then used to transfer the digestion buffers to the S-trap wells. A similar custom protocol on the Opentrons was used to normalize peptide input amounts for StageTip desalting by transferring specified volumes from the S-trap deep-well elution plate to a secondary plate (Bio-Rad, Cat. No HSP9601) and adjusting to a standard volume. StageTip desalting was carried out in a 96-well plate format by removing the base from a used S-trap plate and inserting the StageTips into the wells, allowing for processing using a benchtop centrifuge (Beckman Coulter) and elution into a 96-well plate (Bio-Rad, Cat. No HSP9601). The samples were dried in the elution plate and stored at −80°C until LC-MS/MS analysis.

### BRC & CRC FFPE tissue macrodissection

For BRC and CRC blocks, 10 µm sections were stained with hematoxylin and eosin (H&E) to identify and macrodissect tumor-rich areas that were transferred to a Covaris 96-well plate in quadruplicates (performed at VARI). Each replicate consisted of tumor-rich areas obtained from 5 scrolls equivalent to a total surface area of 30 mm^2^. To evaluate optimal tissue storage conditions, two of the four experimental replicates from each block were stored with or without TLB. The samples went through the S-trap digestion and SDB-RPS StageTip desalting workflows described above. A pooled sample consisting of all BRC & CRC peptide samples was fractionated with SDB-XC StageTips as described above.

### LC-MS analysis of BRC & CRC FFPE samples

For each of the BRC & CRC FFPE samples, 1 µg of peptides (prior to StageTip desalting) were loaded on a Vanquish Neo^TM^ UHPLC system (Thermo Fisher) coupled to an Orbitrap Exploris^TM^ 480 Mass Spectrometer (Thermo Fisher) operating in data-independent acquisition (DIA) mode. Peptides were separated on a 25 cm New Objective PicoFrit^TM^ column that was packed in-house with 1.5 µm C18 beads (Reprosil-Pur, Dr. Maisch). The separation occurred over a 94-minute active gradient at a 200 nL/min flow rate: 1.6% B - 4.8% B in 1 min, 4.8% - 24% B in 83 min, then 24% B - 48% B in 10 min. MS1 spectra covered a mass range of 350-1800 m/z with a resolution of 120,000 at 200 m/z. The MS1 automatic gain control (AGC) target was set to 3E6 with a maximum injection time of 25 ms. For DIA analysis, 30 variable windows were used to cover m/z 500-740 as shown in supplemental Table S2. The scans were performed at an AGC target of 5E5 with a maximum injection time of 54 ms. For fragmentation, higher energy collisional dissociation (HCD) was set to 27% normalized collision energy (NCE) with default charge state set to +3. The DIA method had an average cycle time of 2.2 seconds, equating to ∼5 data points per peak (DPPP). The pooled and fractionated BRC/CRC samples were run on the same LC-MS setup described above, with the instrument operating in data-dependent acquisition (DDA) mode. Data was acquired at a resolution of 60,000 with an AGC target of 3E6 for MS1. MS2 spectra were acquired using a Top-20 paradigm with a resolution of 120,000 and an AGC target of 3E5. Precursors of charge 2-6 were fragmented using HCD at 30% NCE.

### Evaluation of NCOA4 overexpression by TMT or label-free DIA

Four mouse FFPE blocks were obtained from Dr. Joseph D. Mancias at the Dana-Farber Cancer Institute (19). The blocks included embedded tissues from multiple different organs with two blocks being from wild type animals and two from animals with induced overexpression of NCOA4. The blocks were sectioned into four experimental replicates and processed with the optimized Covaris, digestion, and StageTip parameters outlined above.

For DIA analysis, 1 µg of peptides (measured prior to SDB-RPS StageTip) were injected on various LC-MS systems. This analysis encompassed a variable-window DIA method on the Orbitrap Exploris 480, Orbitrap Astral^TM^ DIA with narrow isolation windows (4 m/z, Thermo Fisher), and ion mobility-based DIA (dia-PASEF) on the timsTOF-HT^TM^ (Bruker Daltonics) with a variable window scheme as shown in supplemental Table S3. The wide-window DIA method on the Exploris 480 was consistent with the method described in the previous section, with the exception of a wider mass range (400-850 Th) to enhance peptide depth as shown in supplemental Table S4.

For dia-PASEF, peptides were separated on a nanoElute-2^TM^ (Bruker Daltonics), using a PepSep^TM^ ULTRA C18 column (Bruker Daltonics, 25 cm length/75 µm inner diameter/1.5 µm particle size) and a 30 minute active gradient at a flow rate of 500 nL/min: 3% B - 28% B in 28 min, 28% B - 32% B in 2 min. The dia-PASEF method on the timsTOF HT covered an m/z range of 300-1400, optimized with py-diAID using two ion mobility windows per m/z area (20), yielding a total of 20 dia-PASEF windows and an additional MS1 scan. The IM scan range was set to 0.7 to 1.45 V cm^−2^ with 50 ms ramp and accumulation time, equating to a cycle time of 1.2 s and ∼6 DPPP. Collision energy was synchronized with the IM range, 59 eV at 1/K_0_ = 1.6 Vs cm^−2^ decreased to 20 eV at 1/K_0_ = 0.6 Vs cm^−2^.

For the Astral DIA analysis, peptides were separated using a Vanquish Neo^TM^ UHPLC system (Thermo Fisher) coupled to an Orbitrap Astral^TM^ (Thermo Fisher) with a Aurora Elite^TM^ TS C18 UHPLC column (IonOpticks, 15 cm length/75 µm inner diameter/1.7 µm particle size) at a flow rate of 800 nL/min. The 21 minute active gradient was as follows: 2 %B - 4 % B in 0.5 min, 4%B - 8 %B in 0.6 min, 8%B - 22.5 %B in 12.8 min, 22.5%B - 35%B in 6.9 min, and 35%B - 55%B in 0.4 min. MS1 spectra covered a mass range of 390-980 m/z with a resolution of 240,000 at 200 m/z using the Orbitrap mass analyzer. The MS1 AGC target was set to 5E6 with a maximum injection time of 3 ms, and MS1 spectra were acquired with a timed loop control of 0.6 s. 4 m/z isolation windows spanned the range of m/z 390-900. Maximum injection time was set to 5 ms for MS2 scans, with an AGC target of 2E5. Normalized collision energy was set to 25%.

For TMT analysis, peptides (∼50 ug) were dried down and reconstituted to ∼1 mg/mL in 100 mM TEAB. Peptides were labeled in a 1:5 ratio with TMTpro 18-plex (Thermo Fisher, Cat no. A52045). All channels were fully labeled, and two channels were designated to represent a mixed pool of peptides from all experimental replicates, which would later be excluded from bioinformatic analysis. Following labeling, the 18 channels were quenched with hydroxylamine to a final concentration of 0.3%. The TMT-labeled samples were then pooled and dried down before desalting with SepPak tC18 (Waters, Cat. No. WATO36820). Offline fractionation of TMT-labeled peptides was performed using a high-pH reversed-phase gradient with a ZORBAX^TM^ C18 column (Agilent, 250 mm length/4.6 mM inner diameter/3.5 µm particle size) as described previously (21). The resulting fractions were concatenated into 24 fractions for LC-MS analysis in data-dependent acquisition (DDA) mode. One µg of each fraction was injected on the Thermo Scientific Vanquish Neo UHPLC coupled to a Thermo Scientific Orbitrap Exploris 480 with a 25 cm New Objective PicFrit^TM^ column that was packed in-house with 1.5 µm C18 beads. A 99-minute active gradient at a flow rate of 200 nL/min was used: 1.8% B - 5.4% B in 1 min, 5.4%B - 27% B in 86 min, and 27% B - 54% B in 12 min. MS1 spectra covered a mass range of 350-1800 m/z with a resolution of 60,000. The MS1 AGC target was set to 3E6 with a maximum injection time of 25 ms. The top 20 precursors were isolated, and a monoisotopic peak selection (MIPS) filter was set to peptides, with the “relaxed restrictions when too few precursors were found” toggle turned on. The precursor intensity threshold for selection was 5E3 with a fit threshold of 50% and a window of 1.2 m/z. Precursors of charge states 2-6 were selected for fragmentation with an isolation width of 0.7 m/z, the dynamic exclusion was set to 20 seconds, and NCE was set to 34%. MS2 scans were acquired with a resolution of 45,000, with an AGC target of 3E5, and a maximum injection time of 105 ms.

### Proteome and phosphoproteome analysis of LUAD FFPE blocks

The LUAD FFPE samples were macrodissected by VARI as described above and directly placed into a Covaris 96 AFA-Tube TPX Plate pre-filled with TLB buffer. The samples had varying total tissue area (9 - 63 mm^2^). Covaris plates with dissected tissue were kept at −80C until time of processing with optimal Covaris, digestion, and StageTip parameters as described above. Proteome and phosphoproteome data were collected on the Orbitrap Astral using the same LC setup and 4 m/z isolation windows DIA method described above.

For phosphopeptide enrichment, 50 µg of peptides following S-Trap digestion were desalted using a Sep-Pak^TM^ tC18 96-well plate (Waters, Cat No. 196002320) and eluted in 200 µL of 80% MeCN/0.1% trifluoroacetic acid (TFA). An additional 1% TFA was added prior to phosphopeptide enrichment on the AssayMAP Bravo liquid handling system (Agilent). Fe(III)-NTA cartridges (Agilent, 5 µL) were initially primed with 100 µL of 50% MeCN/0.1%TFA and equilibrated with 50 µL of 80% MeCN/0.1% TFA, followed by sample loading (195 µL, 5 µL/min). The cartridges were then washed with 50 µL of 80% MeCN/0.1%TFA and the phosphorylated peptides eluted with 20 µL of 1% NH4OH directly into a 96-well plate (Bio-Rad, Cat. no HSP9601) containing 2.5 µL of 100% formic acid. Following elution, 20 µL of 100% ACN was added to each sample using a multi-channel pipette. The samples were dried down and stored at −80°C until LC-MS analysis.

### Data analysis

For all experiments, data was searched against the reviewed human proteome database obtained from UniProt, without isoforms (20,462 entries). Contaminants were appended to the FASTA file, and decoys were generated using the reverse sequence approach.

DIA data from the FFPE workflow optimization and macrodissections were searched in library-free mode on FragPipe V20.0 (https://fragpipe.nesvilab.org/). Default parameters of DIA_Speclib_Quant workflow were loaded.

- Peak matching: precursor mass tolerance: 20 ppm, fragment mass tolerance: 20 ppm, calibration and optimization: mass calibration and parameter optimization, Isotope error: 0/1/2. Require precursor was enabled to discard PSMs without any identified parent peaks.
- Protein digestion was set to strict trypsin, with KR cuts at the C-Terminus and maximal cleavages set to 2. Peptide length was set to a minimum of 7 and a maximum of 50. Peptide mass range was kept at 500 to 5000 Da. N-terminal methionine excision was enabled.
- Methionine oxidation and N-terminal acetylation were set as variable modifications, while carbamidomethylation of cysteine was selected as fixed modification.
- An additional 1% run-specific protein FDR filtering was performed during the DIA-NN quantification step.

The DDA data from the pooled and fractionated BRC and CRC samples were analyzed using FragPipe with the localization-aware open-search default parameters in MSFragger and PTM-Shepherd (https://ptmshepherd.nesvilab.org/). For PTM-Shepherd, the extended output option was enabled to obtain additional information for all PSMs.

All NCOA4 overexpression data were searched using DIA-NN both because of its compatibility with all acquired raw file types and to control for any database searching variability. An in-silico digest spectral library was generated in DIA-NN against a mouse FASTA downloaded from UniProt, with unreviewed sequences to include NCOA4 sequences. The fragment ion range was m/z: 100-2000 m/z, and the precursor ion range was set to m/z 200-1600, with charge states 2-4. The peptide length was 7-30 with a maximum number of variable modifications of 3 and 1 missed cleavage. Carbamidomethylation was selected as a fixed modification. N-terminal methionine excision, methionine oxidation and N-terminal acetylation were chosen as variable modifications. TMT data were searched in FragPipe V20 based on the parameters from the TMT16 workflow. The precursor mass tolerance was set to +/− 20 ppm. Peptide length was set to 7-50 with 2 missed cleavages. Carbamidomethylation of cysteines and TMT-tagging of lysine and N-terminal were chosen as static modifications. N-terminal acetylation, methionine oxidation, pyroglutamic acid, pyrocarbamidomethylation of cysteine, proline hydroxylation and N-terminal deamidation were searched as variable modifications. The maximum number of variable modifications per peptide was set to 3. Phosphorylation data was searched in Spectronaut 18 in direct DIA mode. Cysteine carbamidomethylation was set as a fixed modification, while methionine oxidation, N-terminal acetylation and phosphorylation of serine, threonine and tyrosine were set as variable modifications, with a maximum variable modification count of 5. A PTM probability threshold of 0 was used to allow for phosphosites of Class II (sites with only moderate probability of having been accurately localized) and above, with a cutoff filter of ≥ 0.75 being applied post-analysis to determine accurately localized Class I phosphosites (22).

### MSFragger-DIA and Spectronaut instantiation on the Terra platform

DIA data was searched using the MSFragger-DIA workflow, which was added in FragPipe v20.0(17). MSFragger-DIA was implemented as a Google Cloud-based pipeline on the Terra platform (https://app.terra.bio) using a headless configuration to pass commands to a Fragpipe docker. To enable searching of DIA files collected from ThermoFisher instruments, MSConvert (23) was adapted to Terra for parallelized Fragpipe-compatible raw-to-mzML conversion. The default conversion settings of MSConvert are reflected in the standard Terra configuration: MS1 peak picking (centroiding) and removal of consecutive extra zeros in MS1. The Terra-hosted Fragpipe pipeline requires users to input a GUI-generated workflow file, containing parameters for running relevant Fragpipe modules with full functionality. Additional documentation for the Terra MSConvert and Fragpipe workflows is available on GitHub (https://github.com/broadinstitute/PANOPLY) (24). Spectronaut version 19.3 was also implemented as a Terra pipeline with functionality for multiple report generation, customized enzyme database integration, and combination of SNE files.

### Experimental Design and Statistical Rationale

Experimental design and statistical rationale are provided in the individual method sections and also indicated in the result or figure legends corresponding to technology development workflows that were optimized using BRC and CRC blocks. An FDR of 0.01 was set as a cut-off for peptide and protein identifications. For scatter plots in Fig. 3E, fold-change between OE and WT samples were calculated and plotted on the X and Y axes, respectively, and Pearson correlation is indicated in the upper left corner. Differential expression between OE2 and WT2 was calculated using a two-sample t-test, and the volcano plot in Fig. 3E shows −log10 (nominal p-value) and log2 fold-change.. NMF clustering and Gene Set Enrichment Analysis (ssGSEA) pathway analysis of the LUAD MMD samples were performed using the NMF module within Panoply (https://github.com/broadinstitute/PANOPLY). Details have been described before (24–27). Pathways with FDR < 0.05 are indicated with an asterisk. Use of human samples in this this study was part of the approved institutional review board committee at Broad Institute and the CPTAC consortium.

## Result

### Workflow and method optimization

A schematic representation of the FFPE processing workflow is shown in Fig. 1A. Individual 10 µm tissue scrolls are transferred into wells of a Covaris TPX plate, prefilled with Covaris Tissue Lysis Buffer (TLB). Ultrasonication by AFA is applied for tissue deparaffinization, followed by heating the plate to 90°C for protein decrosslinking. A second AFA ultrasonication step ensures thorough tissue homogenization, effectively bypassing the need for either xylene-based deparaffinization and traditional probe-based sonication. This streamlined approach minimizes sample loss and contamination while enhancing processing efficiency and reproducibility.

**Figure 1:**
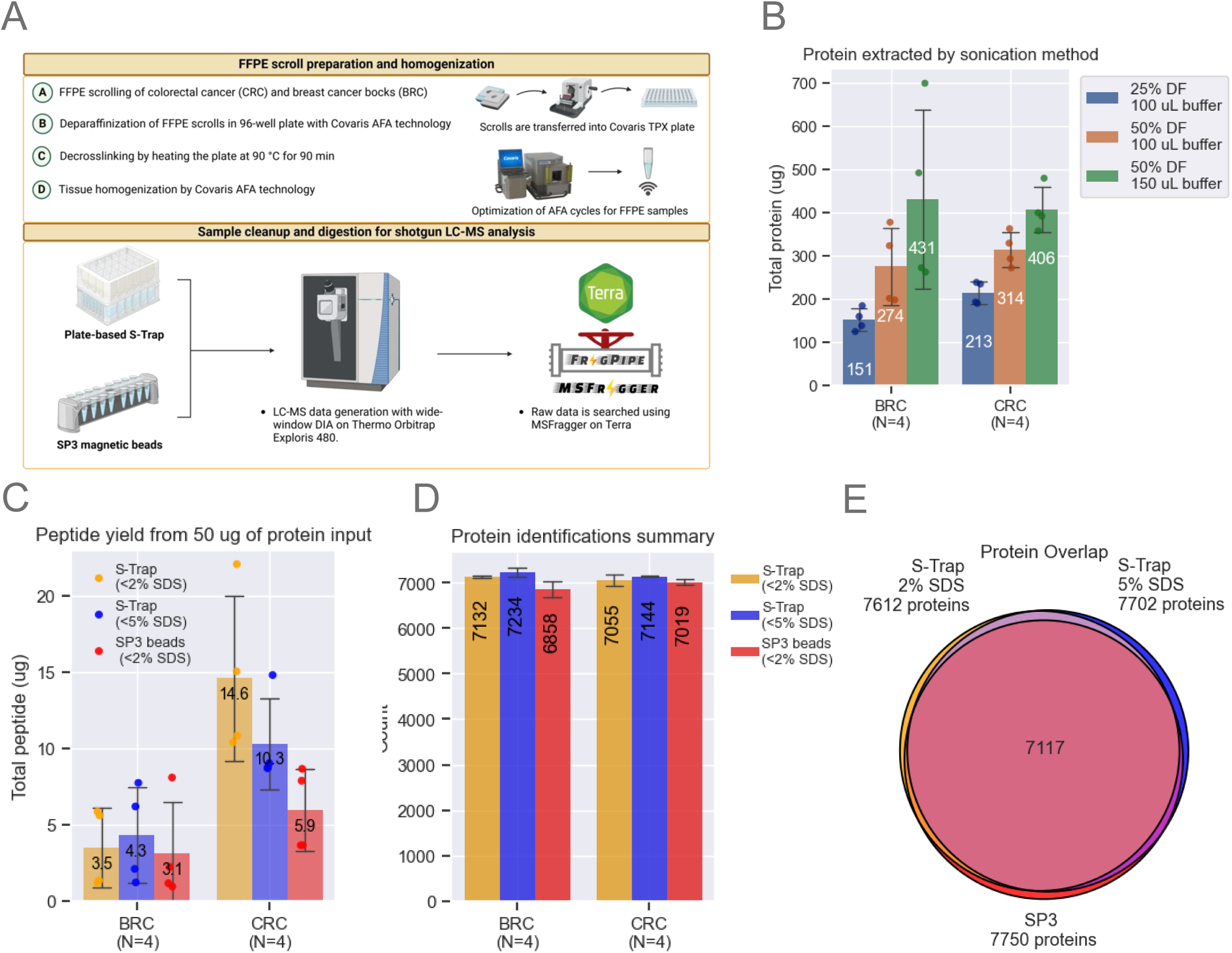
Method Optimization. A. The FFPE workflow processes entire scrolls using a Covaris TPX plate. Breast cancer (BRC) and colorectal cancer (CRC) blocks were scrolled and transferred into a Covaris plate containing tissue lysis buffer. Sonication conditions were optimized to maximize protein extraction. SP3 beads and S-Trap plates were evaluated, starting with 50 µg of protein. Samples were analyzed on a Neo Vanquish LC coupled to an Exploris 480 (Thermo). B. Optimization of protein extraction efficiency by adjusting sonication duty factors (DF) and buffer volume. Bar plots represent data from n=4 scrolls per block, showing means and standard deviation (SD) error bars. C. Comparison of SP3 and S-Trap methods for digestion and cleanup using a fluorometric peptide assay. Bar plots represent data from n=4 scrolls per block, with means and SD error bars. D. Unique protein identifications using MSFragger-DIA across the three tested workflows. Bar plots represent data from n=4 scrolls per block, with means and SD error bars. E. Venn diagram illustrating protein overlap, with each circle representing the total number of unique proteins identified by each tested workflow.

The final workflow developed was the product of systematic optimization of critical sample processing parameters that impact protein/peptide yields, sample cleanup, and the reproducibility of proteomic identifications as detailed below. Our initial experiments focused on refining the recommended Covaris workflow for clinical proteomics (28) using 10 µm scrolls from colorectal cancer (CRC) and breast cancer (BRC) blocks that had been stored ∼20 years (reflecting real-world FFPE repositories). We found that modifications to the tissue lysis buffer (TLB) volume and sonication duty factor (DF) on the Covaris LE220-plus focused-ultrasonicator led to significant improvements in protein yield from the FFPE scrolls. Increasing DF to 50% and TLB volume to 150 µL increased protein yields by more than 100% compared to the Covaris-recommended protocol of 25% DF in 100 µL of TLB (Fig. 1B). A small pellet remained in most wells after processing. Our initial optimization studies utilized the clear lysate (see below).

Following FFPE sample lysis, the protein content of each sample was quantified colorimetrically using the Micro BCA Protein Assay Kit (Thermo Fisher), followed by reduction in 5 mM tris(2 - carboxyethyl)phosphine (TCEP) and alkylation by 50 mM 2-chloroacetamide (CAA). Next, two sample digestion methods, S-Trap (Protifi) and SP3 (Cytiva), the former at two SDS concentrations, were compared with respect to peptide yield starting with 50 µg protein from BRC and CRC FFPE replicate sections (Fig. 1C). Yields from CRC scrolls were ∼3X those from BRC scrolls, possibly due to the higher fat content of the latter, which may interfere with the protein BCA assay, explaining the difference in observed yields across different tissues based on their heterogeneity. BRC peptide yields were roughly comparable across the three tested conditions: S-Trap with tissue lysis buffer (TLB) (∼2% SDS), S-Trap with TLB spiked with additional SDS (∼5% SDS), and SP3 beads with TLB (∼2% SDS). In contrast, peptide yields for CRC scrolls were more than twice as high for S-Trap than for SP3 methods using 2% SDS, and somewhat reduced when S-Trap SDS was increased to 5% (Fig. 1C).

The possible effects of variations in the protocol on the numbers of proteins identified were then evaluated using BRC and CRC FFPE tissue. Comparable numbers of proteins were identified for CRC and BRC samples by analyzing the equivalent of one µg of peptide (normalized based on the peptide yields shown in Fig. 1C) (Fig. 1D, see methods) with over 90% overlap in identified proteins irrespective of the processing method used (Fig. 1E). The S-Trap method, however, identified ∼10% more peptides than the SP3 method (supplemental Fig. S1, supplemental Table S1). Furthermore, total ion chromatograms (TICs) had greater overall intensity and were more reproducible for samples processed by S-Trap than SP3 (supplemental Fig. S1B, supplemental Table S1), despite slightly better digestion efficiency with SP3 (supplemental Fig. S1C, supplemental Table S1). In addition, cleaner samples are obtained using the S-Trap workflow due to an additional chloroform/methanol wash step. Hence all further evaluations utilized S-Trap rather than SP3.

### Application of the optimized method to macro-dissected tissue

The challenges to tissue homogenization and emulsification posed by the wax-encapsulated FFPE tissue samples are ameliorated but not eliminated by powerful sonication devices like the Covaris. Laser microdissection (LMD) of tumor-rich areas is the established strategy for addressing those challenges and pathologist-guided manual macro-dissection (MMD) provides a viable alternative (29). Moreover, enrichment for tumor epithelial cells within heterogenous cancer tissue can aid in inferring cancer-specific proteins, post-translational modifications (PTMs), and pathways.

For a subset of CRC and BRC samples, we optimized parameters for pathology-guided macrodissection of tumor-rich areas from three to five 10 µm thick scrolls per sample (Fig. 2A, supplemental Table S2). Transferring fragments from MMD samples to empty wells was impeded by electrostatic interaction; however, transfer was made easier and more consistent when wells were pre-filled with TLB. To assess the downstream impact of wet vs. dry stabilization, samples transferred to both dry wells and pre-filled wells were frozen at −80°C and processed after several weeks. Protein yields from the clear lysates were nearly identical between MMD samples stored in dry versus pre-filled wells (Fig. 2B), as were peptide yields obtained by processing 50 µg of protein. A single dry-stored BRC2 sample exhibited an unexpectedly high peptide yield (supplemental Fig. S2A). As noted above, despite utilizing a 50% duty factor (DF) and 150 µl TLB, we consistently observed a small pellet in the processed wells. Post-homogenization lysate clarity was inconsistent, as centrifugation never fully removed insoluble pellet particulates. We hypothesized that the unexpectedly higher yield of peptides of this sample might have resulted from our inadvertently loading more of this pellet material into the S-Trap. To examine this possibility, we processed additional BRC1 and CRC1 lysates, this time including the undissolved tissue that we refer to here as “whole tissue” (Fig. 2C). Use of S-Traps enabled the processing of whole tissue rather than clear lysate only owing to the greater cleanliness of fully processed sample. The S-Trap mesh trapped insoluble tissue material, enabling further protein digestion and extraction of additional peptides. After minimizing excess FFPE matrix by MMD processing and incorporating an additional methanol/chloroform S-Trap wash step (see Methods), proteolytic digestion of whole tissue resulted in a greater than two-fold increase in peptide yields (Fig. 2C), while maintaining equivalent depth and protein abundance profile (supplemental Fig. S2B-C). To investigate whether specific proteins were being extracted more efficiently using the whole tissue approach, we calculated relative abundance for each protein within a sample, dividing the abundance of each protein by the protein abundance sum of the sample. The calculated relative abundances of certain cytoskeletal proteins, such as COL1A2 and KRT19, were increased (Fig. 2D). Notably, the relative abundance across all identified proteins was still highly correlated (spearman correlation = 0.92), indicating strong global agreement between clear lysate and whole tissue processing (Fig. 2D). Overall, we identified up to 8,000 unique proteins in each of the MMD samples. An increase of 600 to nearly 1000 in the number of identified proteins (Fig. 2E) relative to the numbers shown in Fig. 1D resulted in part from analyzing 1 µg of peptides rather than the 500 ng used previously. Median protein abundance coefficient of variation (CV) was <20% across 16 MMD samples representing four replicates each from two CRC blocks and two BRC blocks, demonstrating a high level of reproducibility (Fig. 2F).

**Figure 2:**
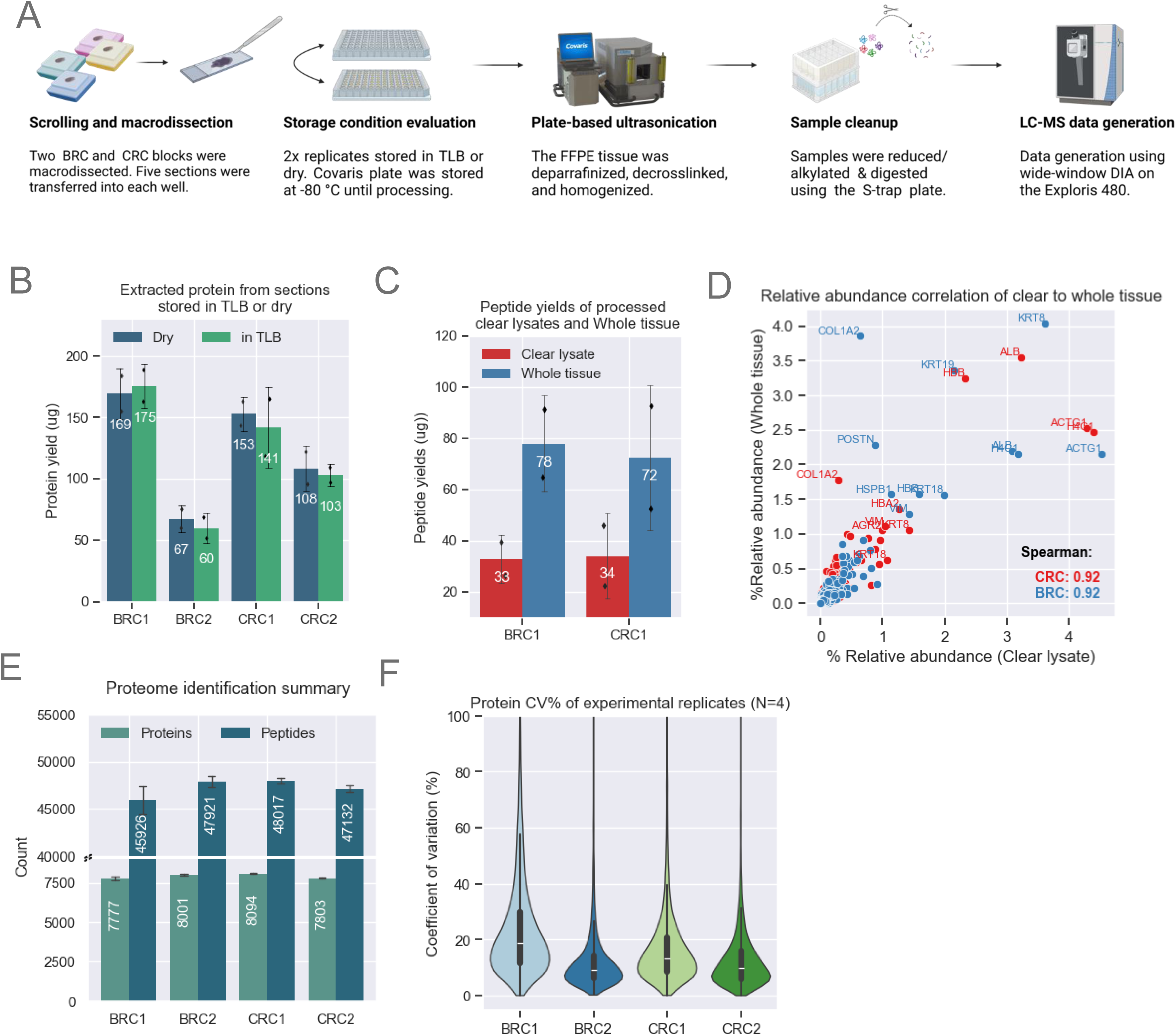
Method Optimization for MMD Samples. A. Schematic showing MMD sample collection and processing. Proteomics data for this figure was generated on an Exploris 480. B. Bar plots showing comparable protein yields (n=2 scrolls per condition) across both storage conditions. The height of each bar represents the mean, with error bars indicating the standard deviation. C. Bar plot showing enhanced peptide yields from whole tissue loaded on the S-Trap compared to clear lysates only. Demonstrated with BRC1 (n=2) and CRC1 (n=2) samples. The height of each bar represents the mean, with error bars indicating the standard deviation. D. Comparison of relative abundance across shared proteins identified from clear lysate and whole tissue. Relative abundance is calculated by dividing each protein’s abundance by the total protein abundance for each sample. Both methods show strong Spearman correlation. E. Unique protein and peptide identifications from MMD tissue with a median area of ∼57 µm² for CRC and BRC samples (n=4). The height of each bar represents the mean, with error bars indicating the standard deviation. F. Protein abundance CVs among experimental replicates (n=4) for each block. The violin plot illustrates data distribution based on the kernel density estimate (KDE). The inner lines represent the 25th, 50th, and 75th percentiles of the IQR range.

However, injecting this amount of sample results in relatively rapid deterioration in column and MS performance. Despite desalting peptides using standard tC18 StageTips, we consistently observed the deposition of white material on the column emitter, likely contributing to performance degradation. To address this, we tested peptide cleanup with styrene-divinylbenzene–reverse-phase sulfonate (SDB-RPS) as the StageTip material, which is known for its potential to improve peptide recovery (30, 31). SDB-RPS cleanup resulted in the near-complete disappearance of these troublesome emitter deposits, enabling LC-MS/MS analysis to be carried out over extended periods without interruption. While no improvements in peptide recovery were observed with SDB-RPS, there was a slight reduction in miscleavage rates. The number of identifications was close to that seen with tC18 (supplemental Fig. S2D-E, supplemental Table S2), with consistency in protein abundance distributions (supplemental Fig. S2F, supplemental Table S2).

Fixation of tissue with formalin results in a range of poorly defined chemical modifications of proteins and peptides (4, 32, 33). The decrosslinking step aims to reverse these formalin-induced chemical modifications. To investigate the decrosslinking efficiency of our workflow, we used FragPipe’s open-search tools to identify fixation-related mass shifts from DDA runs of a pooled BRC/CRC sample by peptide-spectrum matches (PSM) (supplemental Fig. S3A, supplemental Table S2, see methods, (34, 35). The distribution of the observed mass shifts spanned ∼400 Da, with a peak at zero (i.e., no adventitious modification), representing 70% of the PSM library (supplemental Fig. S3B, supplemental Table S2). Approximately 10% of all PSMs exhibited mass shifts, including common formalin modifications such as methylation, formylation, formaldehyde adducts, and methylol groups (supplemental Fig. S3C, supplemental Table S2). These experimental FFPE modifications were largely localized to lysine and arginine residues (supplemental Fig. S3D, supplemental Table S2), consistent with previously reported studies (4). Globally, we identified approximately 74,000 peptidoforms, with 13% of peptides identified only in their chemically modified form, and 25% of peptides showing both modified and unmodified forms (supplemental Fig. S3E, supplemental Table S2). The percentage of chemically modified peptides found in our study is somewhat smaller than previously reported (4).

### Evaluation of quantitative accuracy using multiple instrument platforms

Quantitative accuracy and reproducibility of DIA-generated data can be compromised as a result of fragment ion interference, especially as gradients are shortened. To assess this, we tested instruments that offer high-throughput capabilities, including the Bruker timsTOF HT and the ThermoFisher Orbitrap Astral. The detailed timsTOF HT and Astral data acquisition methods are described in Methods (supplemental Fig. S4A. Supplemental Table S3).

To assess quantification, we utilized FFPE blocks obtained from wild-type (WT) mice and mice over-expressing (OE) NCOA4 (19). As illustrated in Fig. 3A-B, we conducted three independent experiments using scrolls obtained from the two WT and two OE blocks. Each FFPE block contained multiple organs from a single WT or NCOA4-expressing mouse, with a few specific exceptions highlighted in Figs. S4C-D. Aliquots of each sample were analyzed on the Exploris 480 (110 min gradient), timsTOF HT (35 min gradient), and the Orbitrap Astral (24 min gradient). DDA data was also generated by TMT labeling ∼50 ug peptide aliquots of four replicates each of WT and OE in an 18-plex format (supplemental Table S3). Following offline fractionation into 24 fractions, data was acquired on the Exploris 480 using MS2-based peptide quantification. DIA data were searched using DIA-NN on Terra (see Methods), while DDA data was processed using FragPipe.

**Figure 3:**
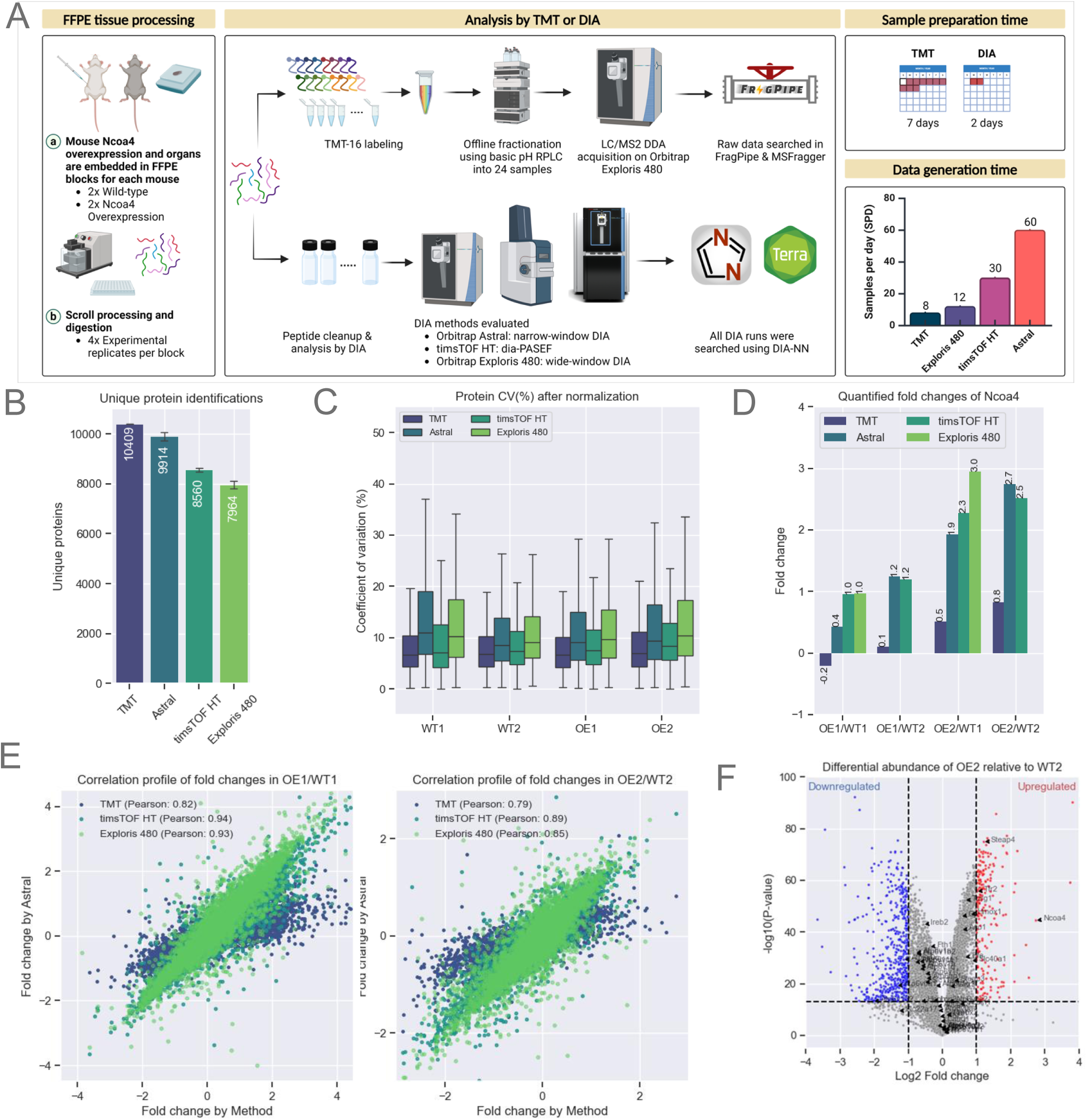
Evaluation of Quantitative Accuracy Using Multiple Instrument Platforms. A. Schematic representation of mouse Ncoa4 WT and OE FFPE blocks. Peptides were either TMT-labeled or analyzed label-free using various DIA methods and instruments, as highlighted in the figure. TMT data was searched using FragPipe while all DIA analysis were performed using DIA-NN as that supported both Thermo and Bruker raw files. B. Protein identifications for each experiment (n=16 per experiment). Bars represent the average, with error bars indicating the standard deviation. C. Normalized protein abundance CVs from experimental replicates (n=4 per sample). The box represents the interquartile range (IQR), with the top and bottom edges indicating the 75th and 25th percentiles, respectively. The line within the box denotes the median, and whiskers extend to include data within 1.5x the IQR. E. Quantified Ncoa4 fold-change between WT and OE mouse models across each TMT and DIA method (X-axis) compared to DIA on Astral (Y-axis). Scatter plots show protein fold-change for two WT and OE pairs, labeled WT1, WT2, OE1, and OE2. F. Volcano plot illustrating differential protein abundance between WT2 and OE2. Significant up or down-regulated (p-value < 0.05) proteins are indicated by red and blue respectively.

As anticipated, the highly fractionated TMT DDA data provided the deepest coverage, encompassing over 10,000 unique proteins. However, the need to analyze each of the 24 fractions using a 110 min gradient resulted in an effective sequencing speed of just 8 samples per day (SPD) (Fig. 3B). In contrast, DIA on the Astral (4 m/z windows) yielded ∼9,900 unique proteins per sample using a 24 min gradient, corresponding to 60 SPD. diaPASEF on the timsTOF HT yielded ∼8,500 unique proteins per sample using a 35 min gradient, corresponding to 30SPD. Wide-window (∼15 m/z) DIA on the Exploris 480 identified ∼8,000 unique proteins using a 110 min gradient corresponding to 12 SPD (Fig. 3B, supplemental Table S3). DDA analysis of the highly fractionated TMT-labeled sample quantified ∼108,000 unique peptides, while Astral DIA achieved a depth of ∼90,000 unique peptides per sample and dia-PASEF ∼74,000 unique peptides per sample (supplemental Fig. S3B, supplemental Table S3). Given the slower speed of the Exploris 480, DIA acquisition on this instrument used wider windows and a shorter mass range compared to the Astral (25 Da windows, m/z: 400-850 vs. 4 Da, m/z: 380-980) and quantified ∼56,000 unique peptides, 40% fewer than the other methods we tested. The median protein abundance CVs for each of the four experimental replicates were all below 20%, calculated using the DIA-NN quantification algorithm in FragPipe without any sample-wise normalization (Fig. 3C).

Differential expression between OE and WT was evaluated to assess quantitative reliability and reproducibility of the TMT and DIA datasets. NCOA4 was prominently up-regulated in the OE FFPE samples in each of the DIA datasets (Fig. 3D), with high correlation of protein fold-change between OE and WT across the different platforms (Pearson’s correlation > 0.8) using the DIA dataset generated on the Astral platform as a comparator (Fig. 4E). As expected, the TMT data were highly compressed relative to the DIA results despite the use of 24 off-line fractions.

**Figure 4:**
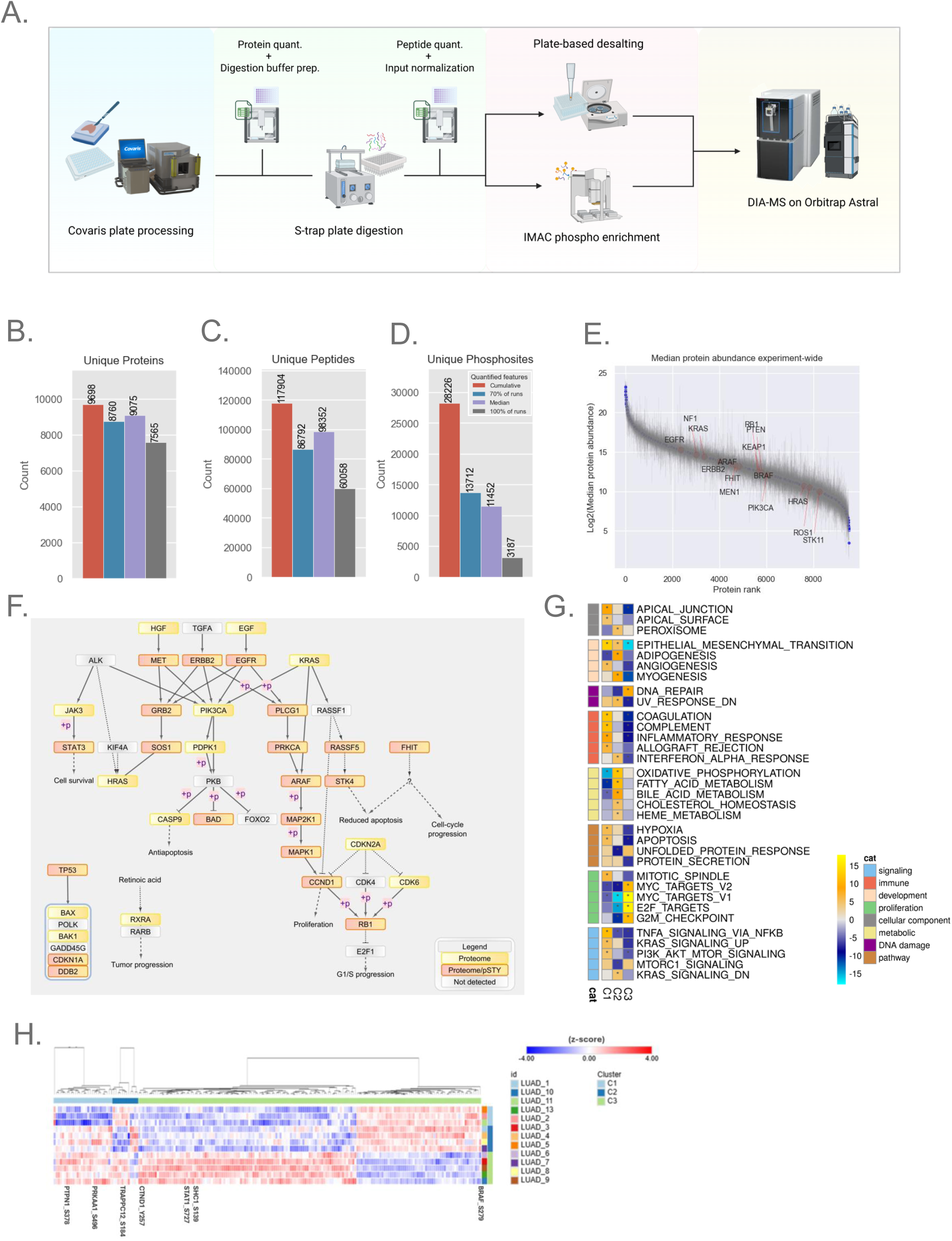
Application to Clinical LUAD FFPE Samples3. A. Semi-automated plate-based workflow for processing of LUAD samples. Opentrons OT-2 automation steps utilized CSV worksheet inputs for easily adjusting plate-to-plate volumes (see Methods for details). B-C. Bar plot showing unique protein and peptide depth from lung adenocarcinoma (LUAD) FFPE blocks processed with MMD (n=13). D. Phosphoproteome depth from LUAD FFPE blocks (n=12). The bar plot represents confidently localized (localization probability > 0.75) and unique phosphosites from 12 samples, each with 50 µg of peptide available for enrichment. E. Rank plot displaying the dynamic range of quantified proteins across all blocks; each point represents median expression, and error bars indicate the quantified range for the experiment. F. Pathway diagram highlighting non-small cell lung cancer pathway-relevant proteins and phosphosites quantified by the workflow. Nodes detected by proteome alone or by both proteome and phosphoproteome are marked with yellow and pink-yellow boxes, respectively. G. NMF clustering of the proteome dataset, resulting in three clusters (C1-3). The heatmap shows pathway terms from Gene Set Enrichment Analysis (GSEA) using the MSigDB hallmark gene sets and pathways with FDR < 0.05 is indicated with an asterisk. H. Heatmap showing the significant phosphosites driving NMF clustering, with a handful of sites highlighted.

NCOA4 selectively regulates autophagic degradation of ferritin, the key cytosolic iron storage complex (36). In addition to observing the expected upregulation of NCOA4 protein, we also observed increased levels of proteins involved in iron metabolism such as Steap4, a metalloreductase that reduces Fe3+ to Fe2+ (37), and Tfr2 (transferrin receptor 2), which allows iron transportation inside cells, likely reflecting feedback mechanisms in response to NCOA4 overexpression (Fig. 3F).

### Semi-automated 96-well plate FFPE proteome and phosphoproteome workflow

Having established a processing pipeline capable of deep-scale proteome profiling of FFPE BRC and CRC tissue, we aimed to assess the feasibility of phosphoproteome analysis and enhance the workflow by incorporating automated components to enable efficient 96-well plate processing (Fig. 4A). Two enabling features of the Opentrons OT-2 robot include the ability to vary digestion buffer volumes for loading on the S-trap plate wells and to normalize peptide input amounts post-digestion, ensuring precision and reproducibility throughout the workflow. By implementing a strategy for StageTip desalting in a 96-well plate format (see Methods), along with plate-based phosphopeptide enrichment and LC-MS injections, the workflow is optimized for high-throughput analysis. The performance of this optimized pipeline was benchmarked using a cohort of lung-adenocarcinoma (LUAD) FFPE blocks of resection tissues from 13 patients. The blocks were macrodissected to enrich tumor-rich sections and digested for global proteome analysis. Twelve of the 13 samples yielded enough peptide to allow enrichment of phosphopeptides from 50 µg input using Fe(III)-NTA AssayMap cartridges on the automated Agilent Bravo liquid handling robot (see Methods).

Based on the robust performance of the Orbitrap Astral in prior experiments, this instrument was selected to evaluate the LUAD FFPE samples (see Methods) using the 60 SPD (24 min) Astral DIA (4 m/z) method. This approach cumulatively quantified approximately 9,700 unique proteins and 120,000 peptides, with a median of 9,075 unique proteins and 98,000 peptides per sample (Fig. 4B-D, Supplemental Table S4). Anticipating that the Astral’s enhanced performance would similarly benefit phosphoproteome analysis, we applied the same Astral DIA method (4 m/z) to these samples. In total, we identified ∼28,000 fully localized unique phosphosites, with a median of 11,000 fully localized phosphosites per sample (Supplemental Table S4). The abundance of quantified proteins spanned a dynamic range of over 20-fold (Fig. 4E).

To verify the biological relevance of quantified proteins, we mapped these proteins back to the non-small cell lung cancer (NSCLC) pathways in the KEGG database. We captured 36 proteins that are actively implicated in the NSCLC cancer pathways reported by KEGG (38), of which 21 also exhibited phosphorylation events (Fig. 4F). Given the limited clinical metadata attached to these samples, we further explored the intrinsic structure of the cohort using non-negative matrix factorization (NMF)-based clustering of proteome or phosphoproteome datasets (26). NMF clustering using proteome and phosphoproteome resulted in three clusters, as supported by multiple metrics including the dispersion and cophenetic correlation coefficients as well as membership consensus (supplemental Fig. S5A-C). Both proteome- and phosphoproteome-only NMF clustering also separated samples into three groups. Some samples clustered differently in the phosphoproteome-only clusters, although overall pathway patterns between the individual clusters were retained (Proteome: Fig. 4F., Phosphoproteome: supplemental Fig. S5E). Phosphoproteome-only clustering may capture subtle heterogeneity in tumors, although the small sample size hindered us from deeper exploration. The first proteomics cluster (C1) showed upregulation of gene sets related to immune signaling such as coagulation, interferon alpha response and proteins related to EMT and apical junction. The second cluster (C2) demonstrated upregulation of metabolic pathways such as oxidative phosphorylation, fatty acid metabolism, and downregulation of Myc and E2F targets. The third cluster (C3) was characterized by downregulation of immune signaling pathways, and upregulation of Myc targets and DNA repair (Fig. 4G). Although the cohort is small, the GSEA results illustrate the ability of standard data analysis approaches applied to FFPE proteomic data to yield biologically relevant molecular characterization of clinical samples. At the level of the phosphoproteome, we identified differential phosphorylation of several key proteins with an established role in LUAD (supplemental Fig. S5F), and explored the phosphosites that contributed to phosphoproteomic NMF clusters (Fig. 4H). In C1, phosphorylation events such as S496 in PRKAA1 and S378 in PTPN1 are relatively downregulated, potentially reflecting a shift in cellular processes such as energy metabolism (39) and phosphatase activity regulation (40). In C2, S184 in TRAPPC12 was relatively downregulated, with dephosphorylation events in TRAPPC12 known to be necessary for cell cycle progression (41). In cluster C3, several notable phosphorylation events were differentially regulated. Phosphorylation of BRAF at S729, a site known to act as a negative feedback regulator, was lower compared to other clusters. Downregulation of the inhibitory site may lead to BRAF-mediated MAPK/ERK signaling, leading to enhanced cellular proliferation. These findings suggest that the oncogenic MAPK/ERK pathway may be more active in C3 (42, 43). Additionally, SHC1 phosphorylation was upregulated at S139, a key site in receptor tyrosine kinase (RTK) signaling reported to be upregulated in clear cell renal cell carcinoma (CCRCC) and downregulated in colon cancer (38), highlighting a diverse role in oncogenic processes. Also of interest was the increased phosphorylation of CTND1 at Y257, a critical modulator of the Ras-MAPK pathway, influencing cellular proliferation, survival, and migration (44). Lastly, STAT1 phosphorylation at S727 was higher in C3 relative to other clusters, a site known to promote AML cell proliferation and differentiation in JAK-STAT pathway-driven neoplasms (45).

In summary, using this small sample set, we demonstrate the overall feasibility of identifying over 9,000 unique proteins from FFPE LUAD tumors and of mapping those proteins to relevant biological processes. The identification of ∼28,000 localized unique phosphosites, with a median of 11,000 phosphosites per sample, further demonstrates the feasibility of phosphopeptide analysis of FFPE tissue samples using our workflow.

## Discussion

We have developed a pipeline comprising high-throughput FFPE tissue processing workflows and optimized data acquisition methods for proteomic and phosphoproteomic applications with increased data depth and completeness as compared to previous DIA-based FFPE analyses. By utilizing ultra-sonication and adaptive focused acoustics, we eliminate the need for xylene-based deparaffinization. Tissue digestion using plate-based S-Trap methods with an additional chloroform/methanol wash and SDB-RPS StageTip desalting allowed for cleaner samples. Label-free DIA analysis shortens the time needed for sample preparation as it avoids chemical labeling, achieving a turnaround time of less than a week for proteome-level analysis of a 96-well plate of FFPE material, with an additional 3 days required for phosphoproteome analysis. The use of DIA in this platform achieves a proteome depth similar to that obtained with TMT, identifying up to 10,000 unique proteins from each sample, but with much faster sample processing and data generation. These same approaches are applicable to tissue microarray samples (manuscript in preparation).

While PTM enrichment in FFPE samples has traditionally been challenging due to sample preservation and processing complexities, our study demonstrates its feasibility in a label-free setting. Using an automated workflow, over 11,000 localized phosphosites were enriched from macrodissected LUAD FFPE tissue samples using the Astral mass spectrometer. Bioinformatic analysis of the proteome and phosphoproteome data generated on a cohort of LUAD samples from archived blocks expands the established utility of proteomics for cancer biology into FFPE sample characterization, with the potential of adding critical insights pertaining to oncogenic mechanisms across multiple omes. The application of proteogenomic data analysis techniques to this small cohort of samples demonstrates the utility of high-throughput, plate-based FFPE proteomics and phosphoproteomics for derivation of meaningful biology from a larger clinical cohort of samples. Our experiments also improve upon the use of DIA for proteomic and phosphoproteomic data collection, allowing for broader applications in proteomics and biomarker discovery of samples. Future studies are aimed at obtaining a deeper understanding of the biological relevance of these and other post-translationally modified peptides and their stability in FFPE samples compared to flash-frozen tissue.

## Author contribution

Conceptualization (MH, MAG, SAC, SS); experimental methodology (MH, JT, MAG, SAC, SS); tissue resources (CN, DCR, GH); formal data analysis (MH, JT, SG); writing, review, editing (all authors).

## Acknowledgements

We also thank Dr. Joseph Mancias and Matthew J Dorman (Dana Farber Cancer Institute, Harvard Medical School) for the NCOA4 wild-type and over-expressing mice FFPE blocks. We also thank Lia Abarzua, Robert Riegelhaupt, Eugenio Daviso and Sameer Vasantgadkar from Covaris, Woburn for their guidance with protocols and Lilian Heil from Thermo Fischer Scientific, San Jose for her assistance with data generation in the Thermo Fischer Science demo laboratory in San Jose, California. Biorender was used to make multiple figure panels in this article (agreement number: SR259LM2NJ). We also thank Natalie Clark and C Williams for assistance with data analysis.

## Funding and additional information

This work was supported by the following awards and grants - Broad Institute SPARC (#800444) awards to S.S and S.A.C, National Cancer Institute (NCI) Clinical Proteomic Tumor Analysis Consortium (CPTAC) grants U24CA270823 to S.S, M.A.G and S.A.C, U01CA271402 to M.A.G and S.A.C and, Dr Miriam and Sheldon G. Adelson Medical Research Foundation to S.A.C.

## Data availability

Data is available on MassIVE (http://massive.ucsd.edu). Username for web access: **[ID not available]**

## Abbreviations

FFPE: Formalin-Fixed Paraffin-Embedded
AFA: Adaptive Focused Acoustics
LC-MS/MS: Liquid Chromatography-Tandem Mass Spectrometry
CVs: Coefficients of Variation
LUAD: Lung Adenocarcinoma
TMT: Tandem Mass Tag
CRC: Colorectal Cancer
BRC: Breast Cancer
TLB: Tissue Lysis Buffer
DF: Duty Factor
BCA: Bicinchoninic Acid
SP3: Solvent Precipitation Beads
TCEP: Tris(2-Carboxyethyl)Phosphine
CAA: 2-Chloroacetamide
MeCN: Acetonitrile
TEAB: Triethylammonium Bicarbonate
FA: Formic Acid
IPA: Isopropyl Alcohol
NH3OH: Ammonium Hydroxide
OT-2: Opentrons OT-2 Robot
H&E: Hematoxylin and Eosin
DIA: Data Independent Acquisition
DPPP: Data Points Per Peak
HCD: Higher Energy Collisional Dissociation
NCE: Normalized Collision Energy
MIPS: Monoisotopic Peak Selection
DDA: Data-Dependent Acquisition
OE: Overexpressing/Overexpression
SPD: Samples Per Day
dia-PASEF: Data-Independent Acquisition - Parallel Accumulation Serial Fragmentation
NSCLC: Non-Small Cell Lung Cancer
KEGG: Kyoto Encyclopedia of Genes and Genomes
NMF: Non-Negative Matrix Factorization
RTK: Receptor Tyrosine Kinase

## Supplementary Figure Legends

**Supplemental Fig. 1:**
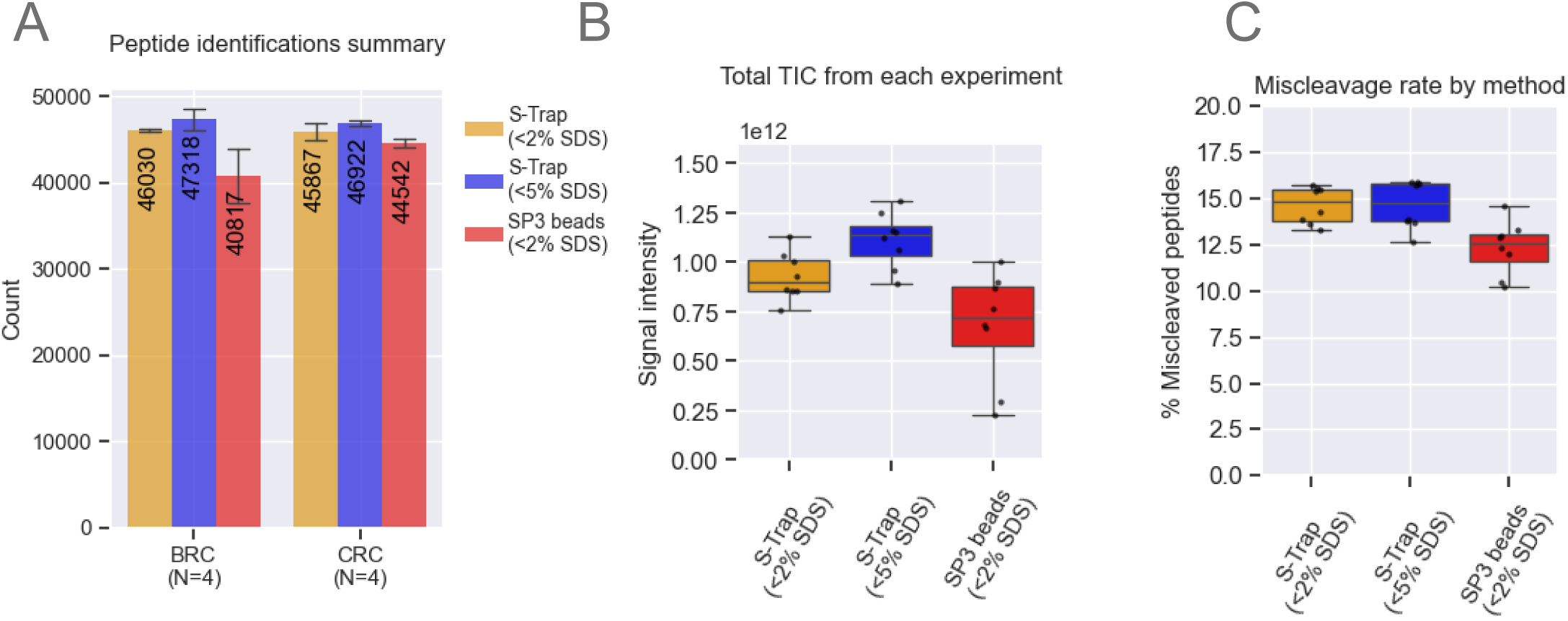
Method Optimization. A. Peptide summary from all workflows. Bar plots represent data from n=4 scrolls per block, showing means and standard deviation (SD) error bars. B. Quantified TIC of all peptides from each experiment across n=4 replicates. The box represents the interquartile range (IQR), with the top and bottom edges indicating the 75th and 25th percentiles, respectively. The line inside the box denotes the median, and whiskers extend to capture data within 1.5x of the IQR. C. Digestion efficiency of S-Trap and SP3 workflows. Bar plots represent data from n=4 scrolls per block, showing means and SD error bars. The box represents the IQR, with the top and bottom edges indicating the 75th and 25th percentiles, respectively. The line inside the box denotes the median, and whiskers extend to capture data within 1.5x of the IQR.

**supplemental Fig. S2:**
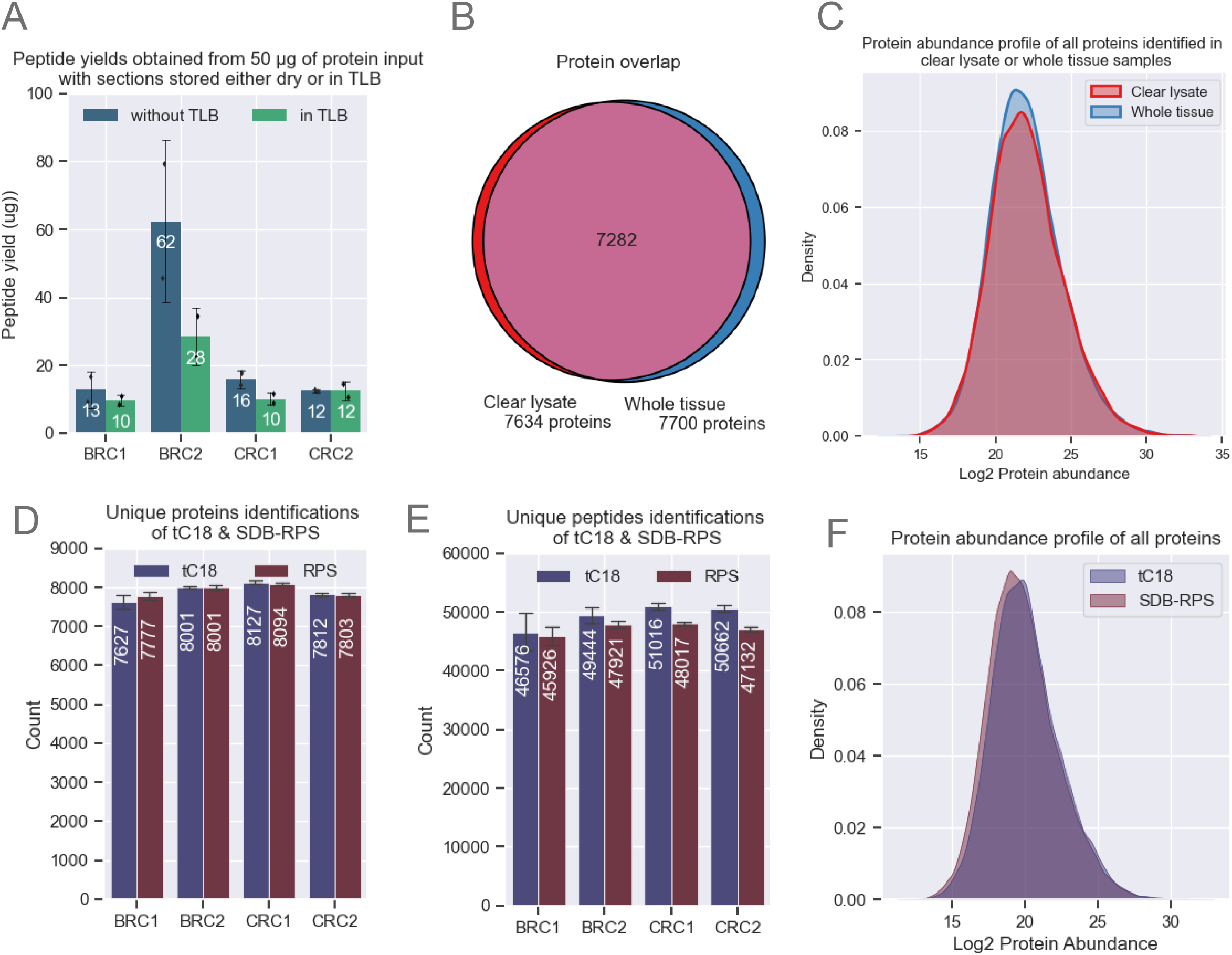
Comparative Evaluation of Sample Collection in Wet vs. Dry Wells and Different Stage-Tipping Materials. A. Peptide yields remain consistent across storage conditions (n=2 scrolls per condition) for each block. The BRC2 block shows the highest yields, likely due to processing the full tissue homogenate, which may increase protein/peptide yields. B. Protein identifications and overlap between whole tissue and clear lysate samples. C. Density plot showing the protein abundance profile, which remains consistent between whole tissue and clear lysates. The peak shape represents the kernel density estimate of all proteins after log2 transformation and normalization. D-E. Comparison of protein and peptide depth in samples desalted with tC18 and SDB-RPS. Each bar represents data from four independent 10 µm scrolls. F. Density plot showing protein abundance for all identified proteins using tC18 and SDB-RPS. The peak shape represents the kernel density estimate of all proteins after log2 transformation.

**Figure S3:**
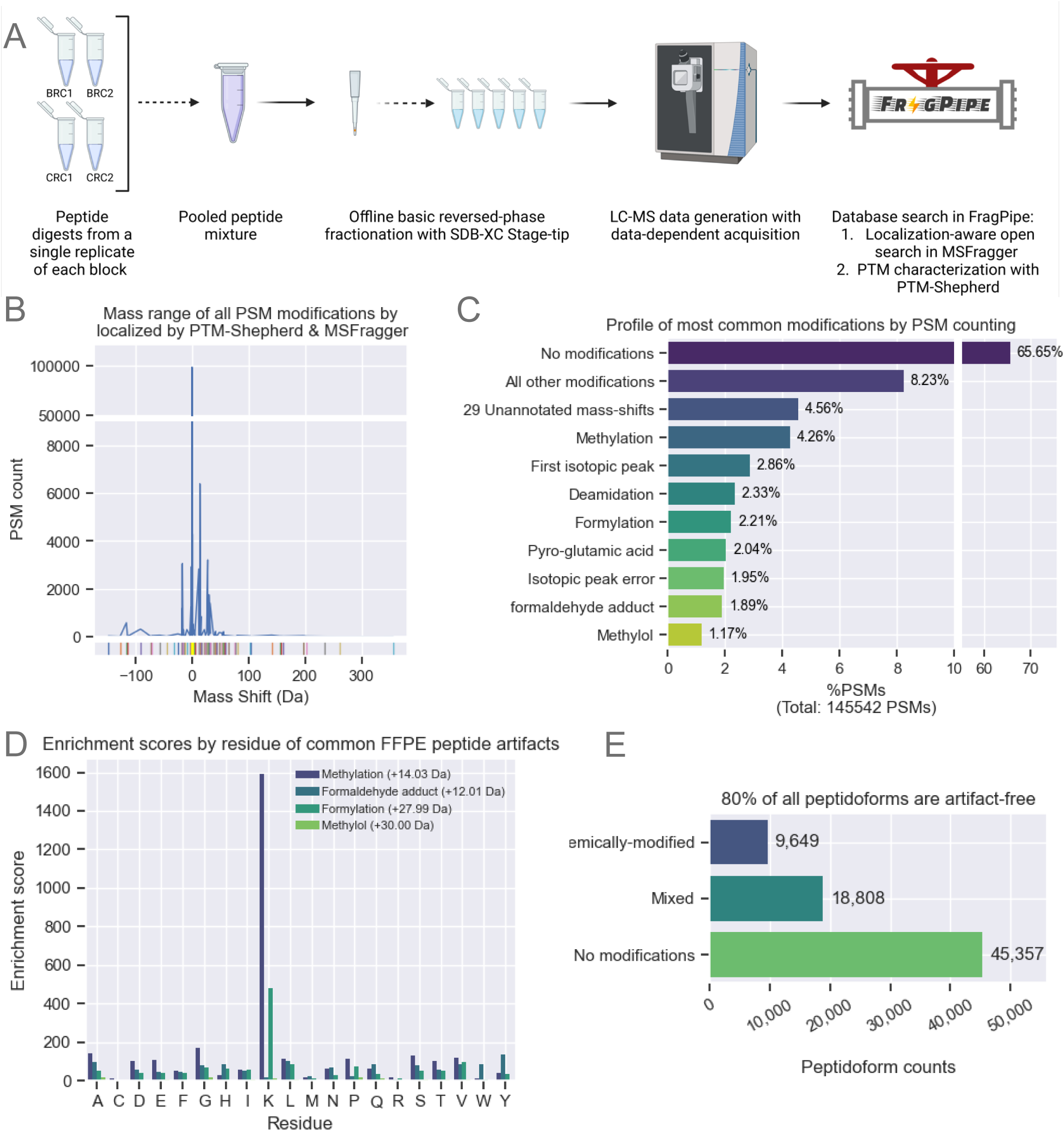
Evaluation of Artifactual FFPE-Derived Peptide Modifications. A. Peptide digests from single replicates of CRC and BRC blocks were pooled equally and fractionated using SDB-XC stage tips into five fractions for data-dependent acquisition (DDA). Data were analyzed using the open-search workflow in FragPipe. B. Open searching with MSFragger and PTM-Shepherd identified 99 different modifications, with most modifications centered around 0 Da. The data were generated from DDA runs of stage-tip fractionated samples pooled from CRC and BRC groups. MSFragger was used for searching, and PTM-Shepherd for modification characterization. C. Peptide Spectrum Match (PSM) summary of all modifications. FFPE-related modifications make up less than 10% of all PSMs. D. Enrichment scores by residue for the most common FFPE modifications. The enrichment score represents the PSM count for each localized residue associated with a specific modification. These scores are representative of the first modified residue in each PSM for a relative enrichment estimate. E. Global modification profile represented by the count of identified peptides from the fractionated pooled sample.

**Figure S4:**
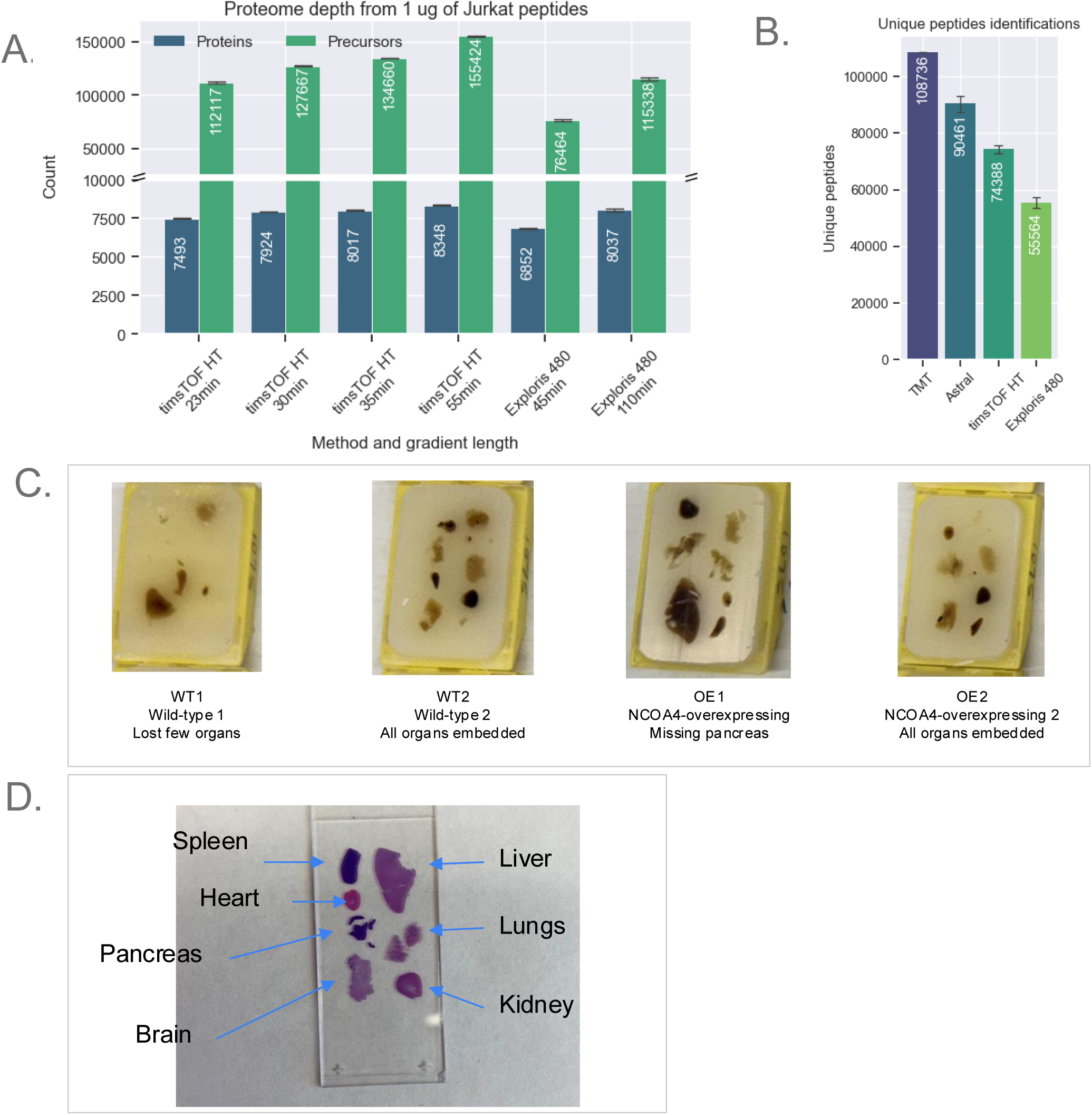
Evaluation of Different DIA Methods and Description of Mouse FFPE Blocks. A. Proteome depth from 1 µg of Jurkat peptides analyzed using optimized DIA methods on the timsTOF HT and Exploris 480. Bars represent the average of (n=2 injections), with error bars showing the standard deviation. The Bruker timsTOF HT, equipped with a dual trapped ion mobility spectrometry (TIMS) tunnels, allows for parallel accumulation-serial fragmentation (PASEF), separating ions by their collisional cross-section (CCS), allowing precursor selection based on their ion mobility (IM) and mass to charge (m/z). Variable-window diaPASEF was optimized for FFPE proteomics using Jurkat peptide digests across four gradients (23, 30, 35, and 55 minutes) on a 25 cm PepSep column. The results were compared to data acquired on the Orbitrap Exploris 480 with 110-minute and 45-minute gradients using a 25 cm home-packed Reprosil C18 column. The wide-window DIA method on the Orbitrap Exploris480 utilized variable isolation windows ranging from 12 m/z to 24 m/z, adjusting to precursor density. Both methods aimed for six data points per peak (DPPP) for quantitative reproducibility. The timsTOF HT 35-minute gradient achieved ∼8,000 unique proteins, comparable to the 110-minute Orbitrap gradient, with a balanced unique peptide depth (∼135,000). B. Peptide identifications across the four methods tested. C. Four blocks from genetically engineered mouse models (GEMM) were used in the mouse FFPE experiments. Two blocks with total body Ncoa4 overexpression (OE) were compared to wild-type (WT) blocks. Blocks WT2 and OE2 included all organs, while WT1 lacked a few organs, and the OE1 block was missing the pancreas. D. Slide representation showing all organs embedded into the GEMM FFPE blocks.

**Figure S5:**
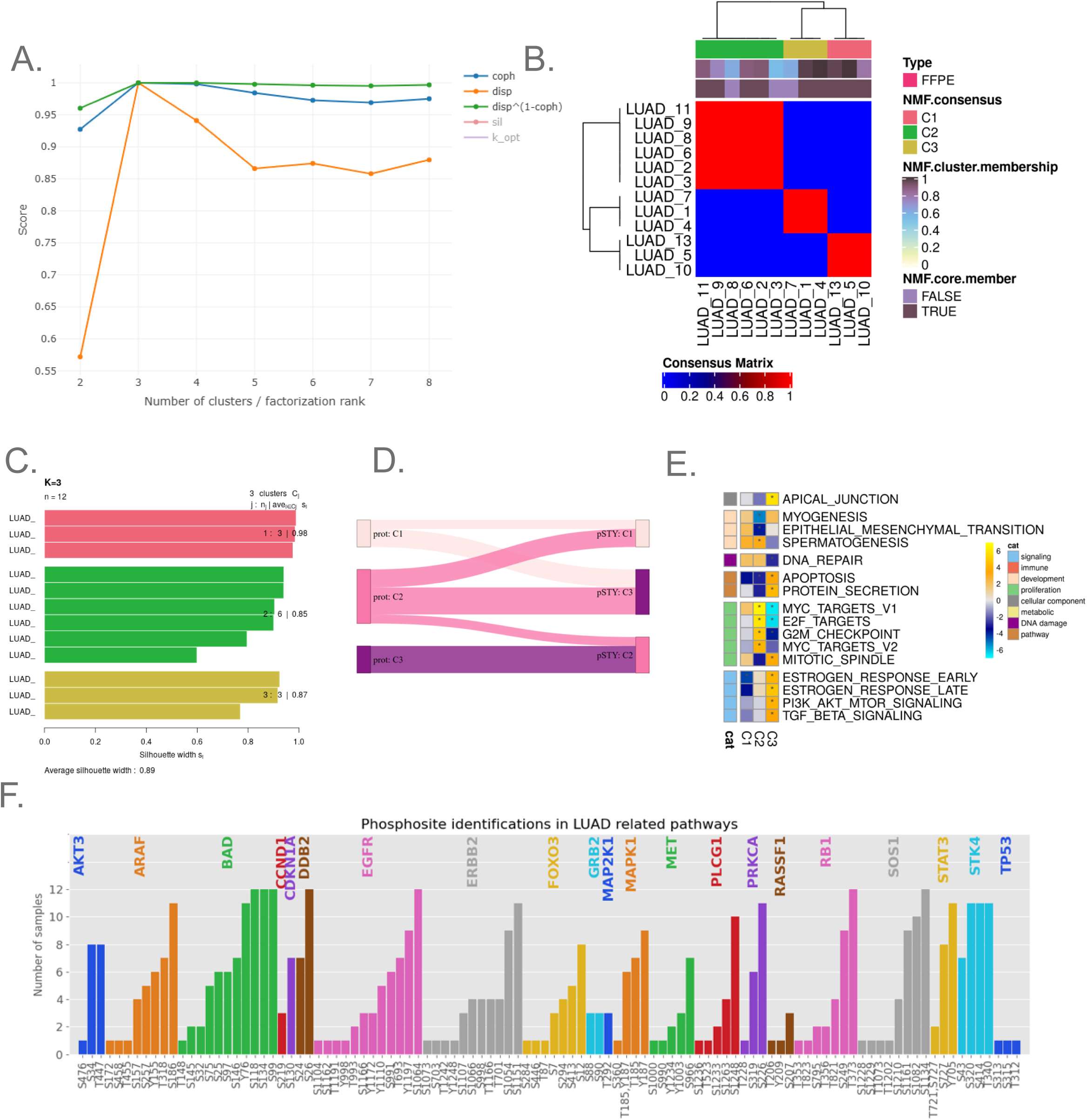
NMF Clustering of Proteome and Phosphoproteome. A. Cophenetic correlation and dispersion scores for each NMF proteome cluster count, with three clusters identified as optimal based on their intrinsic structure and high reproducibility. B. Heatmap showing the clustering stability of LUAD samples into three distinct groups, as defined by NMF consensus and membership scores for each sample. C. Silhouette scores for each LUAD sample based on NMF cluster assignment, used to evaluate the quality of assigned memberships. D. Sankey diagram depicting NMF cluster memberships derived from either proteomic or phosphoproteomic datasets. E. ssGSEA results from the three phosphoproteome NMF clusters. F. Key phosphorylation events and their completeness across the LUAD 12 samples.

## Supplemental Table Legends

**Supplemental Table S1: Global proteomics data associated with development and optimization of the workflow for analysis of the colorectal and breast cancer FFPE blocks**

(A) DIA isolation scheme used for data generation on the Thermo Orbitrap Exploris 480. Method was used to analyze 500 ug of peptides, desalted by tC18 StageTips, derived from breast cancer (BRC) and colorectal cancer (CRC) FFPE blocks. Acquired raw files were searched using MSFragger-DIA Quant workflow.

(B) Protein-level abundance matrix of BRC and CRC blocks processed in 2% SDS and plate-based S-Trap.

(C) Protein-level abundance matrix of BRC and CRC blocks processed in 5% SDS and plate-based S-Trap.

(D) Protein-level abundance matrix of BRC and CRC blocks processed in 2% SDS and SP3 magnetic beads.

(E) S1E: Stats file from the MSFragger-DIA output for BRC and CRC blocks processed by S-Trap (2% SDS or 5% SDS) and by SP3 beads (2% SDS)

**Supplemental Table S2. Tables associated with the method application to macrodissected colorectal and breast cancer FFPE scrolls**

(A) Dimensions of macrodissected tumor sections from breast cancer (BRC) and colorectal cancer (CRC) FFPE blocks, and their protein yields.

(B) DIA isolation scheme used for data generation on the Thermo Orbitrap Exploris 480. Method was used to analyze 1 ug of peptides, desalted by tC18 or SDB-RPS StageTips, derived from breast cancer (BRC) and colorectal cancer (CRC) tumor-rich FFPE sections.

(C) Protein-level abundance matrix from CRC and BRC peptides desalted with tC18StageTips.

(D) Protein-level abundance matrix from CRC and BRC peptides desalted with SDB-RPSStageTips.

(E) PTM-Shepherd summary

**Supplemental Table S3. Global proteomics data associated with the quantitative evaluation across multiple platforms**

(A) DIA isolation scheme used for data generation using the wide-window DIA method on the Thermo Orbitrap Exploris 480.

(B) DIA isolation scheme used for data generation using the diaPASEF method on the Bruker timsTOF HT.

(C) S4C:DIA isolation scheme used for data generation using the 4 m/z DIA method on the Thermo Orbitrap Astral.

(D-G) Protein-level abundance matrix from FFPE blocks, embedding organs originating from genetically-engineered mouse models (GEMM), including wild-type (WT) and Ncoa4 over-expressing (OE) samples, acquired by wide-window DIA on the Thermo Orbitrap Exploris 480 (D), TIms TOF (E), Astral (F), TMT (G) used for data generation using the 4 m/z DIA method on the Thermo Orbitrap Astral.

**Supplemental Table S4. Global proteomics and phosphoproteomics data associated with the application of this workflow to lung adenocarcinoma samples**

(A) DIA isolation scheme used for data generation using the 4 m/z DIA method on the Thermo Orbitrap Astral. The method was used to analyze lung adenocarcinoma (LUAD) tumor rich FFPE sections

(B) Protein-level abundance matrix obtained from LUAD FFPE samples.

(C) Phosphosite-level abundance matrix obtained from LUAD FFPE samples.

(D) Protein-level matrix obtained from non-negative matrix factorization (NMF) clustering of LUAD samples.

(E) Phosphosite-level matrix obtained from NMF clustering of LUAD samples.

